# High-quality human preimplantation embryos stimulate endometrial stromal cell migration via secretion of microRNA hsa-miR-320a

**DOI:** 10.1101/2020.01.21.913632

**Authors:** Robbert P. Berkhout, Remco Keijser, Sjoerd Repping, Cornelis B. Lambalk, Gijs B. Afink, Sebastiaan Mastenbroek, Geert Hamer

**Author notes:** These authors contributed equally to this article (Sebastiaan Mastenbroek, Geert Hamer).

## Abstract

Implantation failure is one of the major success limiting factors in human reproduction. Despite, the mechanisms that determine successful human embryo implantation remain largely unknown. We here show that high-quality human preimplantation embryos secrete soluble signaling factors, including micro RNA (miRNA) hsa-miR-320a, that promote migration of human endometrial stromal cells (hESCs). By using miRNA mimics and inhibitors, we demonstrate that hsa-miR-320a alone can stimulate migration of decidualized hESCs, accurately resembling the response typically triggered only by high-quality embryos. Transcriptome analysis further demonstrated that this effect is very likely mediated via altered expression of genes involved in cell adhesion and cytoskeleton organization. In conclusion, by secreting hsa-miR-320a, high-quality human preimplantation embryos directly influence endometrial stromal cells, most likely to prime the endometrium at the implantation site for successful implantation. Together, our results indicate that hsa-miR-320a may be a promising target to further increase success rates in assisted reproduction.

## Introduction

Human reproduction is characterized by its inefficiency and relatively low success rate. In natural conception, live birth rates per cycle range from 20-30% with a miscarriage rate of 10%, suggesting that most embryos are lost before or right after implantation ^1–3^. Despite advances in reproductive technologies, implantation failure remains a major limiting factor in improving pregnancy rates in IVF/ICSI.

For successful implantation to occur, properly synchronized development of both embryo and endometrium is required. Embryonic development starts when the zygote initiates consecutive cleavage divisions that ultimately result in formation of a morula at day four and a blastocyst at day five of development ^4^. In IVF/ICSI, embryonic development is monitored daily by morphological scoring, including number, size and symmetry of blastomeres, multinucleation and the percentage of fragmentation of blastomeres ^5–7^. Each of these traits contributes to a comprehensive grade of each individual embryo that is predictive for its chances to implant successfully ^7, 8^. Based on these grades, embryos are being selected to be used for transfer into the uterus or cryopreservation ^9^.

In preparation for implantation, also the endometrium undergoes profound changes. After a surge in luteinizing hormone (LH), the endometrium is exposed to rising levels of progesterone leading to decidualization of human endometrial stromal cells (hESCs) ^10^. Decidualization alters the phenotypical characteristics of hESCs, such as its transcriptome and morphology, in preparation for possible implantation of an embryo ^10^. It has been suggested that directed migration of decidualized hESCs is crucial for implantation to succeed, and that altered migration of hESCs may be associated with reproductive failure ^11–13^. Indeed, when comparing the effects of both low and high quality preimplantation embryos, we have recently shown that only embryos of high morphological quality, i.e. with a low percentage of fragmentation, stimulate migration of decidualized hESCs in an *in vitro* migration assay ^14^. In addition, specifically high quality embryos, and not low quality embryos, inhibit migration of non-decidualized hESCs ^14^.

When an embryo enters the uterine cavity, embryo and endometrium are thought to start interacting to initiate implantation ^15^. Before they physically connect, both have been suggested to already secrete a wide variety of signals, the so-called embryo-endometrium crosstalk, which includes hormones, growth factors and cytokines ^16, 17^. After embedding into uterine epithelial and stromal cells, paracrine and cellular signaling has been suggested to further steer the progress of implantation ^18, 19^.

Besides more traditional signaling factors, research has also focused on the roles of more recently discovered factors such as microRNAs (miRNAs)^20^. These 20-25 nucleotide small non-coding RNA molecules play important signaling roles in various mechanisms of health and disease, such as angiogenesis and carcinogenesis ^21–23^. Biogenesis of miRNAs starts in the nucleus where precursor miRNAs are being transcribed. Following transfer to the cytoplasm, precursor miRNAs are spliced by Dicer into single-stranded mature miRNAs ^24, 25^. Next, these mature miRNAs are assembled into RNA-induced silencing complexes (RISC) that can bind to the 3’-UTR of an analogue target mRNA, leading to translational repression and mRNA decay ^24^.

Preimplantation embryos have been shown to secrete miRNAs during multiple steps of development and implantation in human and various other mammalian species ^26–32^. Several studies have also reported that embryos secrete specific miRNAs that may be associated with the method of fertilization, the underlying infertility diagnosis, their chromosomal status or the pregnancy outcome ^28–32^. However, no clear link between embryo-derived miRNAs and a possible biological mechanism of action during implantation has been described. In the present study, we identify a human embryo-derived miRNA that influences migration of decidualized hESCs during implantation.

## Results

### The migratory effect of embryo conditioned medium (ECM) from high-quality embryos can be abolished by RNase

We first confirmed that, in correspondence with our previous study ^14^, decidualized hESCs migration was stimulated by embryo conditioned medium (ECM) from high-quality embryos compared to empty control medium (p=0.002, Figure 1a, b). To determine whether this stimulatory signal was caused by secreted RNA molecules, we repeated these migration assays but with the addition of RNase. We found that the addition of RNase completely abolished the pro-migratory effect of ECM from high-quality embryos on decidualized hESCs (p=0.008, Figure 1a, b).

**Fig. 1.**
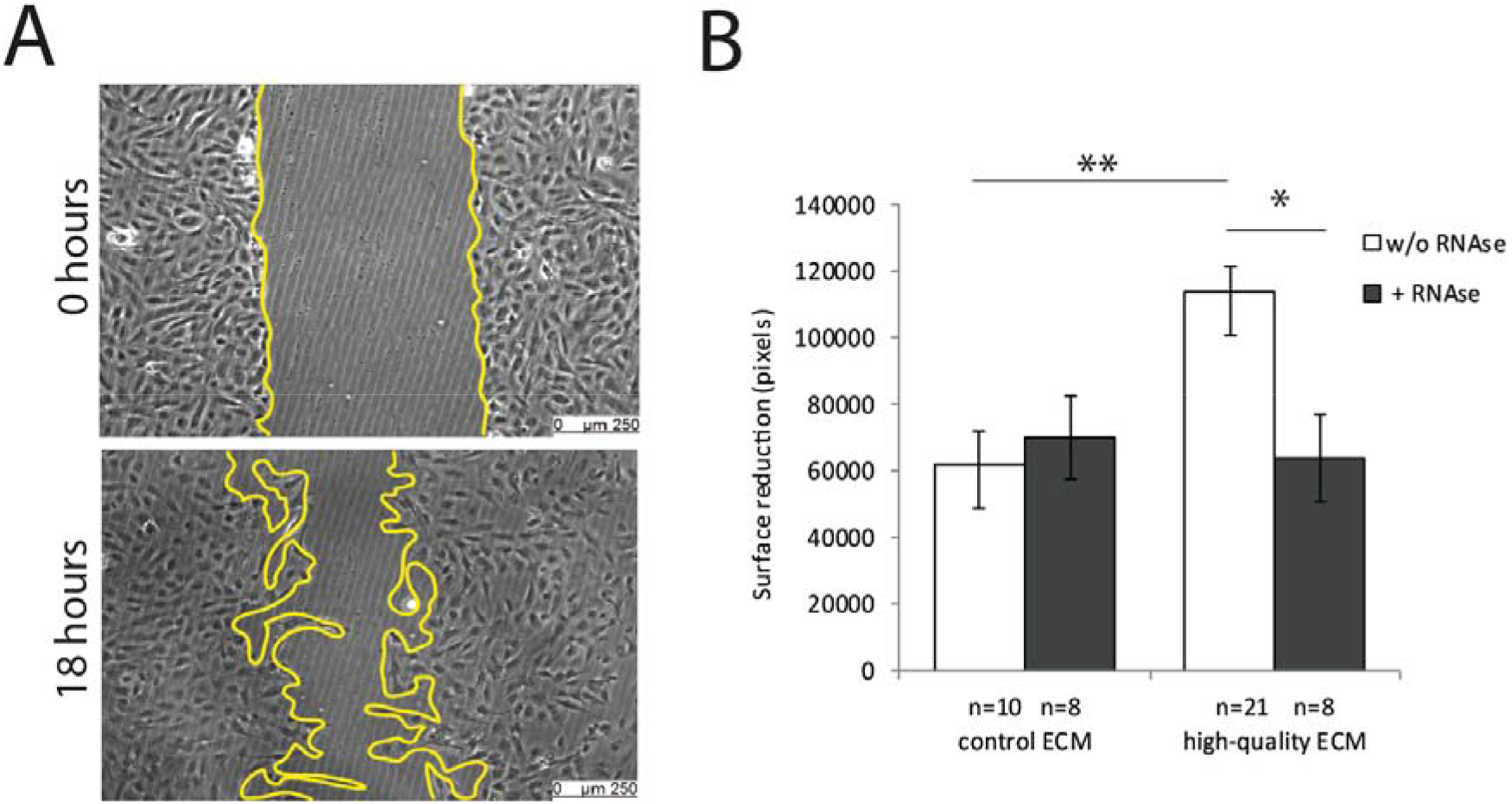
High-quality human preimplantation embryos stimulated decidualized hESCs migration through paracrine RNA signaling. **a-b** Decidualized hESCs migration was stimulated by culture in high-quality ECM compared to control. **b** Stimulatory effect of high-quality ECM on migration of decidualized hESCs was antagonized by simultaneously adding RNase. hESCs, human endometrial stromal cells; w/o, without. Data are expressed as mean ± sem. Values that are significantly different are labelled by asterisks (*p<0.05).

### High- and low-quality human preimplantation embryos secrete specific subsets of miRNAs

To identify the nature of the soluble RNA molecules that stimulated migration of decidualized hESCs, we performed a PCR-array analyzing the presence of 752 human miRNAs in ECM. ECM was collected and pooled based on the morphological quality of the embryos: high-quality embryos (ECM droplets from 48 individually cultured embryos; fragmentation ≤20%), low-quality embryos (ECM droplets from 48 individually cultured embryos; fragmentation >20%) or empty control medium (48 droplets that were prepared and cultured similar to the other groups, but without an embryo being present) (Figure 2a). From each set of 48 ECM samples of either high- or low-quality, 16 samples each were pooled resulting in three biological replicates for both high- and low-quality ECM (Figure 2a).

**Fig. 2.**
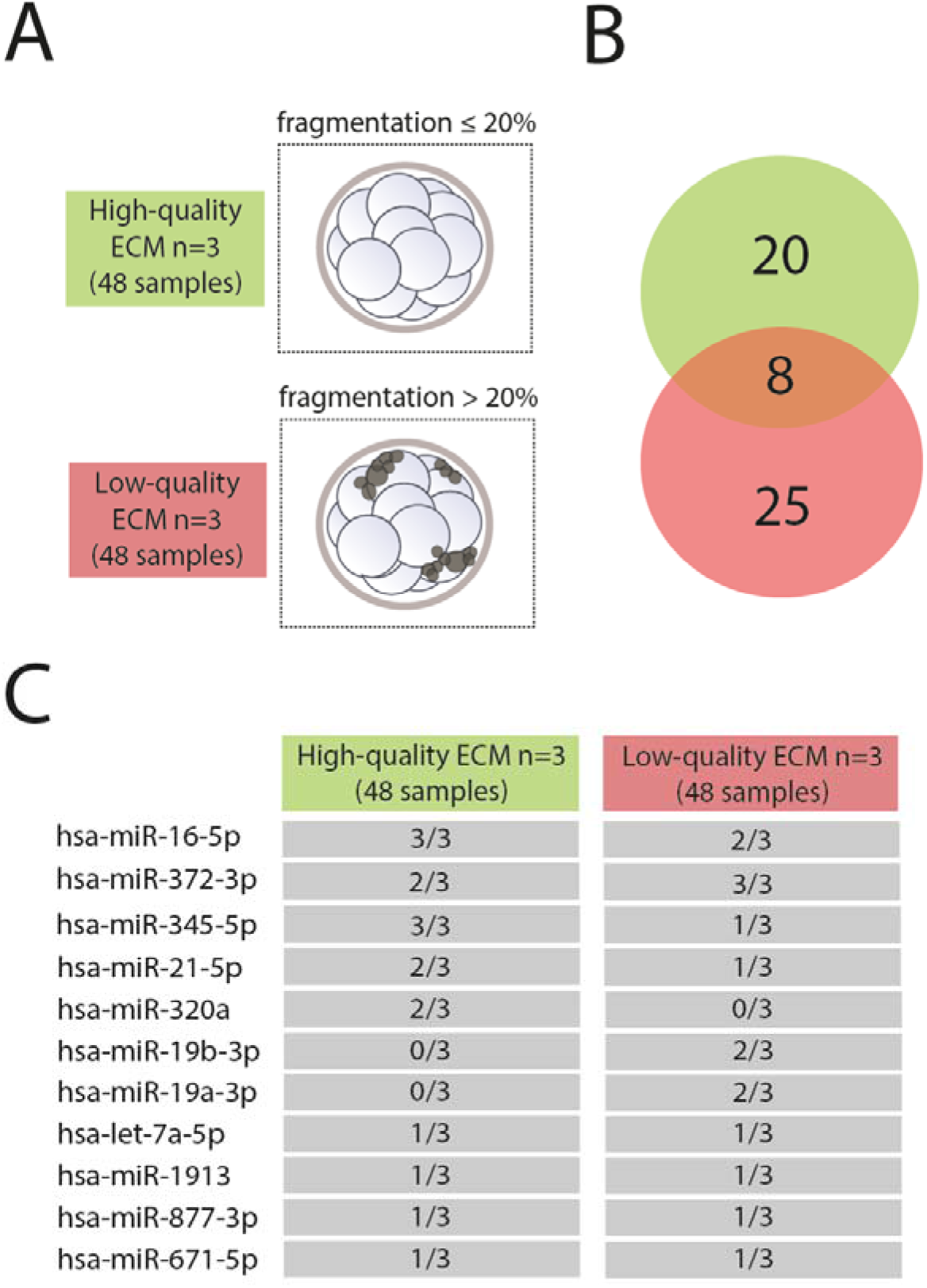
High- and low-quality human preimplantation embryos secrete specific subsets of miRNAs. **a** For both high- and low-quality embryos, 48 individually cultured samples were collected that were pooled per 16 to create 3 biological replicates. **b** High-quality embryos secreted 20 unique miRNAs, low-quality embryos secreted 25 unique miRNAs and 8 miRNAs were overlapping, **c** The most frequently expressed miRNAs were present in both high- and low-quality ECM, except for hsa-miR-320a which is high-quality ECM specific, and hsa-miR-19a-3p and hsa-miR-19b-3p which are low-quality ECM specific. ECM, embryo conditioned medium.

In total, 53 unique human miRNAs were identified in at least one biological replicate, that were not present in control medium: 20 miRNAs unique to high-quality embryos, 25 miRNAs unique to low-quality embryos, and 8 miRNAs that were overlapping between high- and low-quality embryos (Figure 2b). The miRNAs that were most frequently detected in ECM, regardless of the experimental group, were: hsa-miR-16-5p (5/6 experiments), hsa-miR-372-3p (5/6 experiments), has-miR-345-5p (4/6 experiments) and hsa-miR-21-5p (3/6 experiments) (Figure 2c). MiRNAs that were detected in 2/6 experiments were: hsa-miR-320a, hsa-miR-19b-3p, hsa-miR-19a-3p, hsa-let-7a-5p, hsa-miR-1913, hsa-miR-877-3p and hsa-miR-671-5p (Figure 2c). We then identified which miRNAs were exclusively detectable in ECM from either high- or low-quality embryos, and found that hsa-miR-320a was specific to high-quality ECM (2/3 experiments), while hsa-miR-19b-3p and hsa-miR-19a-3p were specific to low-quality ECM (2/3 experiments) (Figure 2c). All other miRNAs that were specific for either high-quality or low-quality embryos were only expressed in one biological replicate. Additional family members of the aforementioned miRNAs were not detected in any of the samples.

### Endometrial stromal cell migration is regulated by miRNA hsa-miR-320a secretion from high-quality embryos

To study the effects of miRNAs that were exclusively secreted by either high-quality or low-quality embryos on decidualized hESCs migration, we transfected hESCs with mimics or inhibitors for either hsa-mir-320a (high-quality ECM) or hsa-miR19b-3p and hsa-miR-19a-3p (low-quality ECM) (Figure 3a). Both mimics and inhibitors were successfully transfected into decidualized hESCs, as visualized by a green fluorescent signal for those miRNAs that could be designed with a fluorescently labelled 5’-FAM tag (Figure 3b). No significant difference in transfection efficiency, determined by the percentage of fluorescently labeled hESCs, was detected between negative controls and miRNA-specific mimics (68.2% vs. 57.6%, p=0.422) or inhibitors (64.7% vs. 53.1%, p=0.100) (Figure 3c).

**Fig. 3.**
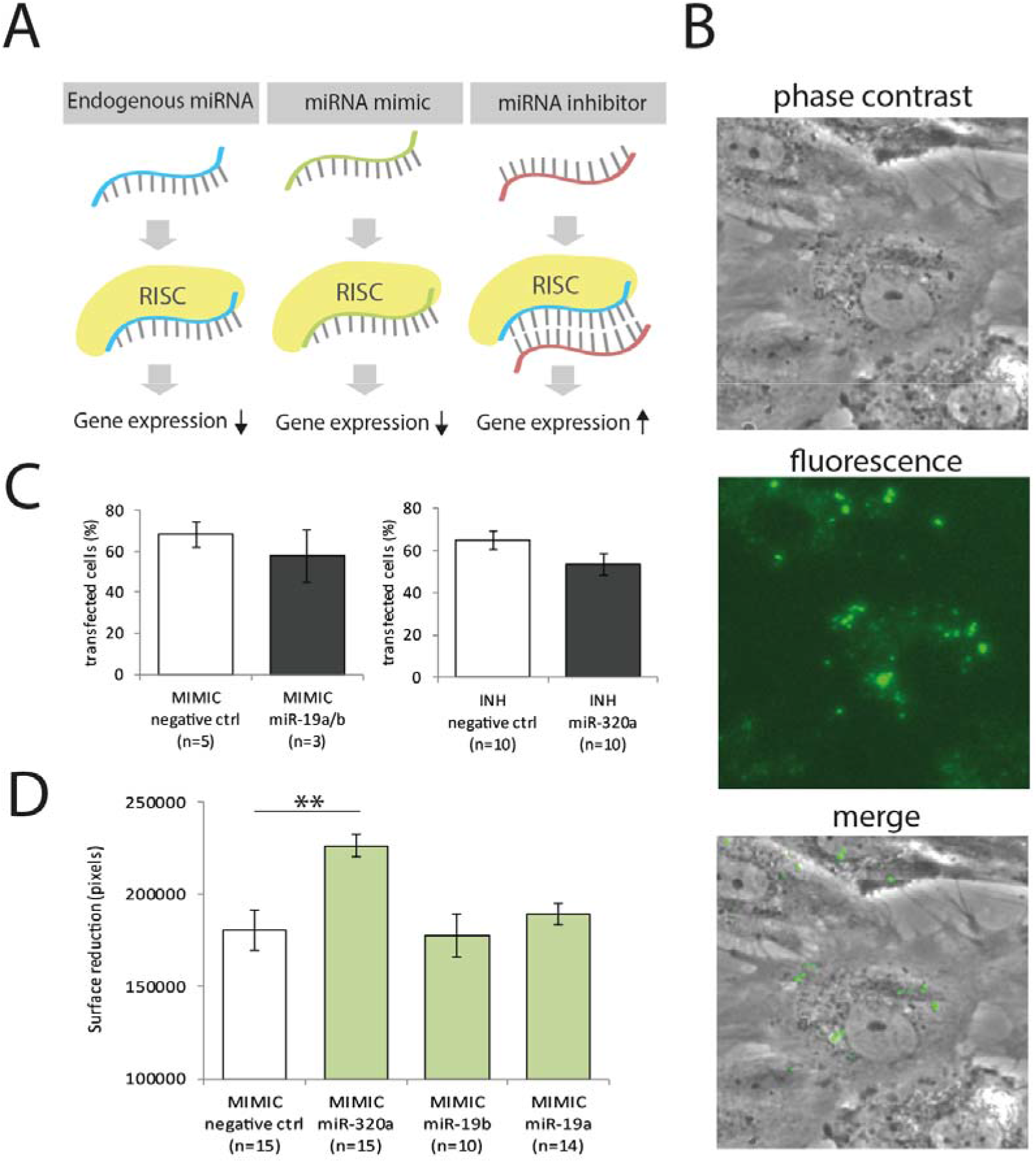
Hsa-miR-320a mimics high-quality ECM-induced migration of decidualized hESCs. **a** Endogenous miRNA are assembled into RNA-induced silencing complexes (RISCs) and suppress expression of target genes; miRNA mimics are likewise assembled into RISCs and suppress expression of target genes; miRNA inhibitors bind to complementary mature miRNA and thereby inhibit miRNA function. **b-c** Transfection percentages of miRNA mimics and inhibitors into decidualized hESCs is not significantly different compared to negative control mimics or inhibitors. **d** Mimics for hsa-miR-320a stimulated migration of decidualized hESCs whereas all other mimics had no effect. hESCs, human endometrial stromal cells. Data are expressed as mean ± sem. Values that are significantly different are labelled by asterisks (*p<0.05).

Next, we identified which of the detected miRNAs was able to mimic ECM induced changes in decidualized hESCs migration, in the absence of ECM. To do so, we first transfected miRNAs into monolayers of decidualized hESCs in the presence of regular culture medium alone. We found that hsa-miR-320a stimulated migration of decidualized hESCs in the absence of ECM (p=0.002), while there was no effect from hsa-miR19b-3p (p=0.995) or hsa-miR-19a-3p (p=0.837), compared to transfection with negative control mimics (Figure 3d). These negative controls were designed to have no homology to any known miRNA or mRNA in human, and are therefore reflective for baseline conditions.

We then investigated whether transfection with hsa-miR-320a mimics would increase the migration response of decidualized hESCs that were subsequently cultured in ECM, derived from low-quality embryos (<8 blastomeres, >20% fragmentation) or empty control medium. To do so, decidualized hESCs were first transfected with hsa-miR-320a mimics, after which migration assays were performed with ECM (Figure 4a). Indeed, decidualized hESCs that were transfected with hsa-miR-320a mimics and that were subsequently exposed to control ECM or low-quality ECM, showed an increased migration response, compared to transfection with negative control mimics (control ECM, p=0.028; low-quality ECM, p=0.040) (Figure 4b).

**Fig. 4.**
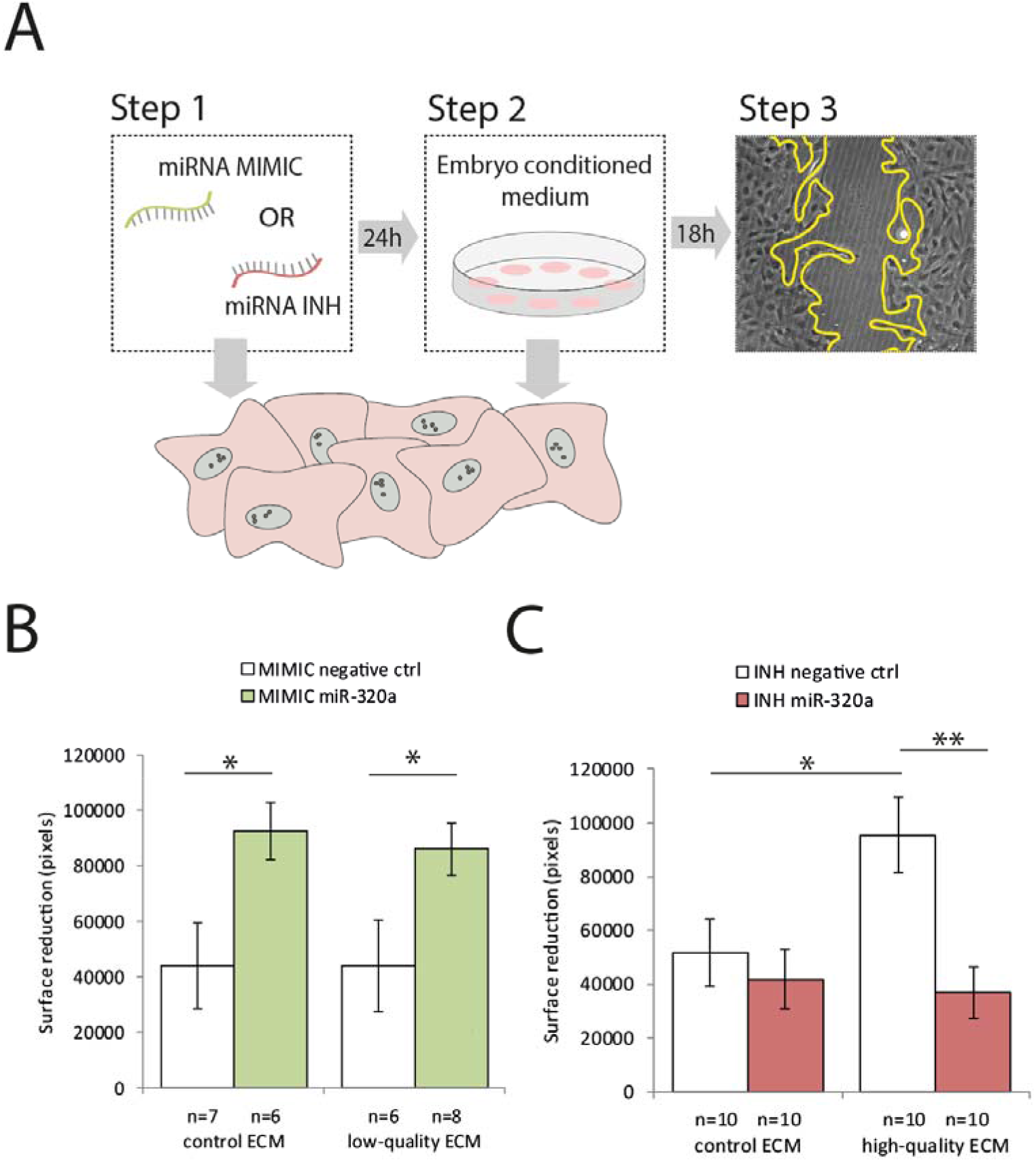
Hsa-miR-320a mimics and inhibitors interfere with ECM to regulate migration of decidualized hESCs. **a** Decidualized hESCs were first transfected with miRNA mimics or inhibitors for 24 hours, after which migration assays (18 hours) were performed during co-culture with ECM. **b** Mimics for hsa-miR-320a stimulated decidualized hESCs migration when co-cultured with control ECM or low-quality ECM. **c** Inhibitors for hsa-miR-320a neutralized the pro-migratory effect of high-quality ECM. hESCs, human endometrial stromal cells. Data are expressed as mean ± sem. Values that are significantly different are labelled by asterisks (*p<0.05).

Next, we examined whether transfection of decidualized hESCs with hsa-miR-320a inhibitors would interfere with signals present in high-quality ECM (≥8 blastomeres, ≤20% fragmentation). First, we confirmed that high-quality ECM stimulated migration of decidualized hESCs after transfection with negative control inhibitors (p=0.034) (Figure 4c). Again, these negative controls were designed to have no homology to any known miRNA or mRNA in human, and are therefore reflective for baseline conditions. Then, after transfection with hsa-miR-320a inhibitors, we demonstrated that decidualized hESCs migration was no longer stimulated by high-quality ECM (p=0.003; compared to control p=0.732) (Figure 4c). In other words, inhibition of hsa-miR-320a completely antagonized any pro-migratory effects of high-quality ECM.

We then questioned whether decidualization was required for hESCs to respond to hsa-miR-320a. To do so, we transfected non-decidualized hESCs with mimics and inhibitors for hsa-miR-320a and repeated the migration assays during co-culture with high-quality ECM (≥8 blastomeres, ≤20% fragmentation) or empty control medium. We found that mimicking or inhibiting hsa-miR-320a function in non-decidualized hESCs did not affect the response to high-quality ECM or empty control medium (Supplementary Figure 1).

### Hsa-miR-320a may influence decidualized hESCs migration by targeting cell adhesion and cytoskeleton organization

Finally, we investigated which specific genes were targeted by embryo-derived hsa-miR-320a in hESCs. To do so, we performed RNA-sequencing on decidualized hESCs derived from 2 patients that were treated with either hsa-miR-320a mimics (n=3 per patient) or negative controls (n=3 per patient). Principle component analysis demonstrated a clear effect of hsa-miR-320a mimics on the transcriptome landscape of decidualized hESCs (Figure 5a). In total, hsa-miR-320a mimics induced differential expression of 223 transcripts. Of these transcripts, 140 were downregulated, and 83 were upregulated in decidualized hESCs treated with hsa-miR-320a mimics compared to negative controls (Figure 5b, supplementary data 1A). Next, we used gene ontology analysis to estimate which biological processes in hESCs could be affected by hsa-miR-320a ^33, 34^. By doing so, we identified various processes that were significantly affected by hsa-miR-320a, of which e.g. cell adhesion, regulation of cytoskeleton organization and cell migration could be directly related to the observed migration response of decidualized hESCs in response to high-quality ECM (False discovery rate of < 0.05) (Table 1, Supplementary Data 1B).

**Fig. 5.**
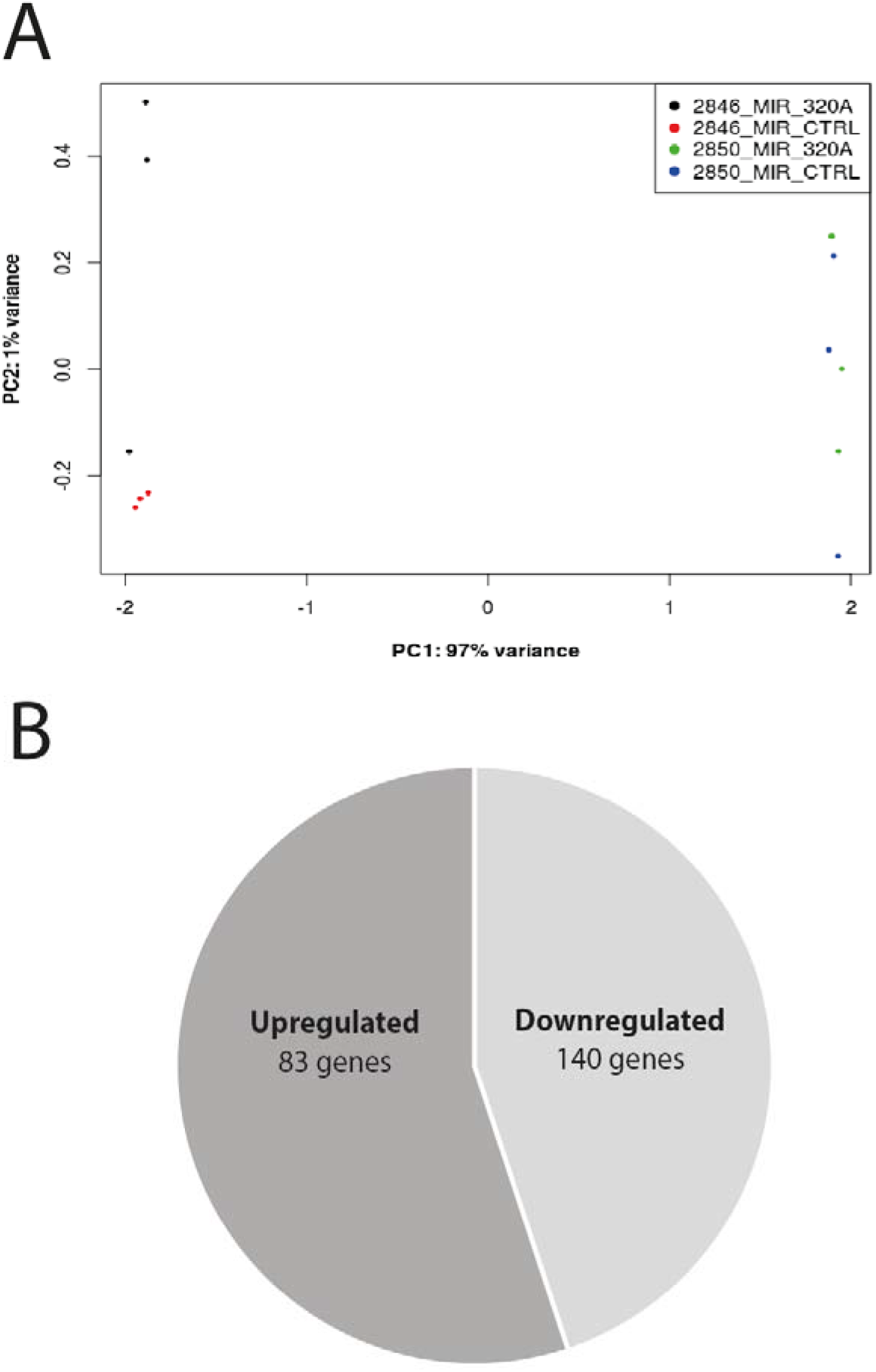
miRNA hsa-miR-320a causes transcriptional alterations in decidualized hESCs. **a** Transcriptional changes in decidualized hESCs from 2 separate patients (2846 versus 2850) are clustered by patient type and transfection type in a multidimensional scaling plot. **b** Upon transfection of decidualized hESCs with hsa-miR-320a mimics or negative control mimics 83 genes were upregulated and 140 genes were downregulated.

**Table 1.** Hsa-miR-320a may influence decidualized hESCs migration by targeting cytoskeleton organization. Gene ontology (GO) analysis, in which enrichment equals −log10(p), where 1.3 is equivalent to p=0.05 and p represents the geometric mean of p-values in an annotation cluster. Only a description of the first term of each statistically significant (enrichment⍰>⍰1.3) annotation cluster is shown. Full results are shown in corresponding supplementary data (Supplementary data 1B).

| Representative GO-term | Description | Enrichment |
| --- | --- | --- |
| GO:0043229 | Intracellular organelle | 4.1 |
| GO:0097190 | apoptotic signaling pathway | 3.5 |
| GO:0044446 | Intracellular organelle part | 3.2 |
| GO:0050839 | Cell adhesion | 3.1 |
| GO:0071840 | Cellular component biogenesis | 2.8 |
| GO:0048514 | Vascular development | 2.6 |
| GO:0009653 | Cell development | 2.5 |
| GO:0009966 | Signal transduction | 2.3 |
| GO:2001233 | Extrinsic apoptosis | 2.2 |
| GO:0005783 | Endoplasmic reticulum | 2.2 |
| GO:0001525 | Angiogenesis | 2.1 |
| GO:1902911 | Kinase activity | 2.1 |
| GO:0005178 | Integrin binding | 2.0 |
| GO:0005178 | Integrin signaling | 1.9 |
| GO:0070273 | Phospholipid binding | 1.8 |
| GO:0007049 | Cell cycle regulation | 1.8 |
| GO:0048523 | Cell signaling | 1.7 |
| GO:0000902 | Cell morphology | 1.6 |
| GO:0051493 | Cytoskeleton organization | 1.5 |
| GO:0010812 | Cell-substrate adhesion | 1.5 |
| GO:0016032 | Viral interaction | 1.5 |
| GO:0048013 | Receptor signaling | 1.4 |
| GO:0005794 | Golgi apparatus | 1.4 |
| GO:1902235 | Endoplasmic reticulum stress signaling | 1.4 |
| GO:0016020 | Membrane components | 1.4 |
| GO:0098552 | Cell surface | 1.4 |
| GO:0031410 | Intracellular vesicles | 1.3 |
| GO:0098805 | Membranes | 1.3 |
| GO:0051270 | Cell migration | 1.3 |

## Discussion

Our work provides evidence that high-quality human preimplantation embryos secrete a specific paracrine signal, miRNA hsa-miR-320a, that drives the endometrial response during implantation. Among 752 miRNAs investigated, hsa-miR-320a was the only miRNA that is specifically secreted by high-quality embryos. Our results further suggest that hsa-miR-320a stimulates decidualized hESCs migration by regulating cytoskeleton organization, a crucial biological process involved in cellular motility^35^.

Migration of various cancerous cells has been previously reported to be affected by miRNAs, including hsa-miR-320a. Interestingly, hsa-miR-320a was most often identified as a tumor suppressor that reduced cellular migration ^36–47^. However, also cases in which hsa-miR-320a stimulated or had no effect on cellular migration have been described ^48, 49^. This discrepancy in reported literature may be attributed to the wide variety of different cell types that have been investigated previously, and potential side effects of hsa-miR-320a on other cellular mechanisms such as proliferation or apoptosis. In addition, the exact molecular mechanism by which hsa-miR-320a acts to regulate cellular motility remains to be clarified. Certainly, our results indicate that organization of the cellular cytoskeleton may be targeted, possibly involving microfilaments, intermediate filaments or microtubules ^35^.

It was reported previously that hsa-miR-320a, which we identified as being secreted specifically by high-quality embryos, as well as miR-19a-3p and miR-19b-3p, which we identified as being specific to low-quality embryos, are being secreted by human blastocysts ^28, 30^. Here, we demonstrate that these miRNAs are already secreted by embryos from as early as the cleavage stage. It was recently shown that hsa-miR-320a is already secreted by oocytes, particularly in those that developed into embryos with high morphological quality ^50^. Moreover, it was shown that hsa-miR-320a functions in an autocrine manner to improve morphological development of preimplantation embryos ^50^. However, the observed effects were contributed to hsa-miR-320 in general, without specification of the exact family member involved. Of note, many miRNAs exist as members of a larger family of miRNAs that have highly similar nucleotide sequences. Although speculative, research has suggested that miRNAs with great sequential similarity, for example members of the same family, may be functionally redundant ^51, 52^. Therefore, considering hsa-miR-320 in general, these studies may together point towards two important aspects of embryo-derived hsa-miR-320. First, presence of hsa-miR-320 distinguishes high-quality oocytes and embryos from their low-quality counterparts. And second, hsa-miR-320 seems to play an important role in autocrine and paracrine signaling by first stimulating proper morphological development and later successful implantation.

Any biomarker that reflects embryo quality in IVF/ICSI may hold great promise for future clinical applications. In this context, miRNA hsa-miR-320a, whose secretion we demonstrate to be specific to high-quality preimplantation embryos, may be suitable for further validation of its predictive value in selecting embryos with the highest potential to implant for transfer. Additionally, artificial supplementation of hsa-miR-320a during embryo culture, to improve morphological development, or during embryo transfer, to improve implantation rates, may be an interesting avenue for future research. In this study we focused on pools of ECM from embryos that had been removed from the culture dishes for cryopreservation and focused solely on hESCs migration as a measure for implantation. Obviously, the perspective of future clinical application requires further research into hsa-miR-320a and its downstream effects on embryo development and implantation.

In conclusion, we identified a soluble miRNA, hsa-miR-320a, that not only marks high-quality human preimplantation embryos, but also directly stimulates decidualized hESCs migration *in vitro*. This makes hsa-miR-320a a unique and promising target for future research to ultimately improve pregnancy rates in IVF/ICSI.

## Materials and Methods

### Ethical approval and informed consent

Regarding the use of endometrial biopsies, this study was approved by the Medical Review Ethics Committee of the Amsterdam University Medical Centers and written informed consent was obtained from all subjects who agreed to donate endometrial biopsies for research purposes. ECM was collected from patients who did not object to storage and use of medical waste tissues and was stored anonymously. According to Dutch law, the use of ECM for this research was allowed without ethical permission.

### Endometrial biopsies

Biopsies were performed on hysterectomy specimens from patients that received surgery for spotting due to a niche in a caesarean section scar (n=2). Patients were pre-menopausal, had a history of proven fertility and at least one live birth, and received no hormonal treatment 3 months prior to surgery. Biopsies were performed randomly in the menstrual cycle and biopsied material was immediately suspended in Dulbecco’s Modified Eagle Medium/Nutrient Mixture F-12 phenol red-free medium (DMEM/F12; L-glutamine, HEPES; Thermo Fisher Scientific, Waltham, MA, USA) supplemented with 1% penicillin/streptomycin at 37°C. Next, the biopsied material was cut into small pieces and further digested enzymatically in DMEM/F12 phenol red-free medium supplemented with >125 U/ml collagenase IV and 1% penicillin/streptomycin. The suspended material was filtered through a 76 μm sterile filter and subsequently cultured in DMEM/F12 phenol red-free medium supplemented with 10% heat-inactivated fetal calf serum (FCS; Thermo Fisher Scientific, Waltham, MA, USA) and 1% penicillin/streptomycin (C-medium). The medium was refreshed after 3 hours to discard all non-attached cells and retain the attached hESCs. Purity of hESCs cultures was assessed by immunostaining for vimentin (M0725, Agilent, CA, USA) and cytokeratin 18 (M7010, Agilent, CA, USA). Only hESCs cultures that were passaged less than 6 times were used for subsequent experiments.

### Embryo conditioned medium (ECM)

All assigned ECM was collected from regular ICSI-cycles in which routine ICSI procedures were followed and during which all embryos were handled according to local regulations and standard operating procedures. Embryos were cultured in Sage medium (Quinn’s advantage cleavage medium, Cooper Surgical, USA) supplemented with 5% human serum albumin (HSA; Vitrolife, Göteborg, Sweden) from day 1-3 after fertilization. Thereafter, embryos were transferred to fresh Sage medium (Quinn’s advantage blastocyst medium, Cooper Surgical, USA) supplemented with 5% HSA, on day 3 after fertilization. Embryo morphology was assessed daily by an independent laboratory technician. Droplets (20 μl) from individually cultured embryos, now referred to as ECM, were collected in sterile 1.5 ml collection tubes after all embryos had been removed from the culture dishes for cryopreservation on day 4. All ECM samples were collected on day 4. Empty medium droplets, in which no embryo was grown but that otherwise underwent the same procedures, from the same dishes were used as controls. Samples were snap-frozen in liquid nitrogen and stored at −80 °C.

### Migration assays

#### Experimental groups

ECM was used from individually cultured embryos with high morphological quality (≥8 blastomeres, fragmentation ≤20%), low morphological quality (≤7 blastomeres, fragmentation >20%) or empty control medium in which no embryo had been cultured, but that otherwise underwent the same laboratory procedures. Per experimental group, ECM from 5 individually cultured embryos was pooled (90 μl).

#### Experimental procedures

30,000 thawed hESCs were cultured in single wells of a 48-wells plate and decidualized by adding DMEM/F12 phenol red-free medium supplemented with 10% heat-inactivated charcoal stripped FCS (Thermo Fisher Scientific, Waltham, MA, USA), 0.5 mM 8-bromoadenosine 3□,5□-cyclic monophosphate (Sigma-Aldrich, Saint Louis, MO, USA), 1 μM medroxyprogesterone acetate (Sigma-Aldrich, Saint Louis, MO, USA) and 1% penicillin/streptomycin (D-medium) for 5 consecutive days. Monolayers of hESCs were scratched with a p200 pipet tip to create a migration zone. HESCs were then cultured in 90 μL of pooled ECM derived from 5 individual embryos. Cultures were overlaid with 150 μL mineral oil (Irvine Scientific, Santa Ana, CA, USA) to prevent evaporation. For addition of RNase, we used 1 mg/ml of RNase in PBS (10109169001, Roche, Basel, Switzerland). At 0 hours and 18 hours after creation of the migration zone, phase contrast images were taken using a monochrome digital camera (Leica DFC365 FX) attached to an inverted microscope (Olympus IX71) with a U Plan FL 4x/0.13 PhL objective. The migration response of hESCs was quantified by assessing the reduction of the surface area of the migration zone by using Image J software (version 1.50i).

### miRNome PCR

#### Experimental groups

ECM from individually cultured embryos of high-quality (fragmentation ≤20%; 48 samples), low-quality (fragmentation >20%; 48 samples) or empty control medium, in which no embryo had been cultured (48 samples), was used. Per experimental group, three biological replicates were performed by pooling ECM samples from 16 individually cultured embryos (320 μl pooled ECM). To ascertain that cellular fragmentation was the only difference among these experimental groups, samples from embryos with different developmental stages (ranging from 6 blastomeres to the morula stage) were evenly distributed across both experimental groups.

#### Experimental procedures

All procedures for miRNome analysis were performed at Qiagen, Hilden, Germany. Total RNA was extracted using miRCURY RNA isolation kit – Biofluids (300112, Exiqon, Denmark) according to manufacturer’s instructions and RNA was eluted in 50 μl RNase-free H2O. Next, RNA was polyadenylated and cDNA was synthesized according to manufacturer’s instructions (miRCURY LNA RT Kit, 339340, Qiagen, Hilden, Germany). Expression of miRNAs was determined by Ready-to-Use Human panel I+II PCR, using ExiLENT SYBR® Green master mix (339322, Qiagen, Hilden, Germany). Amplification reactions were performed in a LightCycler 480 Real-Time PCR System (05015243001, Roche, Basel, Switzerland) in 384 well plates. Amplification curves were analyzed using LC software (04994884001, Roche, Basel, Switzerland), both for determination of Cq values and for melting curve analysis. Only samples with Cq<37 were included in the analysis.

### miRNA transfections

Hsa-miR mimics (339173, miRCURY LNA miRNA Mimic) and inhibitors (339121, miRCURY LNA miRNA Inhibitors) were ordered from Qiagen, Hilden, Germany. Additionally, negative control mimics (YM00479902, Negative Control miRCURY LNA miRNA Mimic) and negative control inhibitors (YI00199006, Negative control A), which both have no homology to any known miRNA or mRNA in human, were ordered from Qiagen, Hilden, Germany. The following miRNAs were labeled with a fluorescent 5’-FAM tag: hsa-miR-19a mimic + inhibitor, hsa-miR-19b mimic + inhibitor, negative control A mimic, negative control inhibitor. 30,000 decidualized or non-decidualized hESCs were cultured in single wells of a 48 wells plate and transfected with 25 nM miRNA mimic or negative control mimic. Alternatively, 30,000 hESCs were transfected with 150 nM miRNA inhibitor or negative control inhibitor. All transfections were performed in D- or C-medium by using HiPerFect Transfection Reagent according to manufacturer’s instructions (301704, Qiagen, Hilden, Germany). After transfection for 24 hours, hESCs were washed with PBS and transfection efficiency was determined by calculating the percentage of fluorescently labelled hESCs from the total number of hESCs per image.

### RNA-seq

#### RNA isolation

Per well of a 6-wells plate 300,000 hESCs were cultured and decidualized for 5 consecutive days, after which hESCs were transfected for 24 hours with 25 nM hsa-miR-320a mimic or 25 nM negative control mimic. Directly after, transfection medium was removed and hESCs were harvested by using a cell scraper (3010, Corning, NY, USA) and passed through a 20 gauge syringe needle. Next, total RNA was isolated using the ISOLATE II RNA Mini Kit (BIO-52072, Bioline, London, UK) according to manufacturer’s instructions.

#### Library construction and RNA-sequencing

Total RNA was sent to BGI Tech Solutions (Hong Kong, China) for library construction and sequencing (BGISeq500, paired end 100□bp). Twelve libraries were constructed for non-stranded pair-end cDNA sequencing on the BGISEQ-500 RNA-Seq platform. Cleaned sequence reads were mapped against the reference human Hg38 build (USC Genome Browser) using HISAT2. The counts of the aligned reads were summarized with featureCounts in the Rsubread package. Multiple mapped reads were excluded from counting. Fold changes were calculated, and differential analysis was performed with the R packages edgeR-Voom. Genes exhibiting differential expression between MIR_320A and MIR_CTRL transfected hESCs with p < 0.05 were included in further analyses.

### Statistical analysis

Mean differences among experimental groups were tested by one-way ANOVA or student’s t-test. To correct for multiple testing we used Dunnett and Tukey post-hoc tests when applicable. In all cases, IBM SPSS Statistics 23 software was used. P-values <0.05 were considered statistically significant.

## Data availability

All sequence data have been submitted to the Sequence Read Archive (NCBI) and are available under the accession number PRJNA601880.

## Author contributions

S.R., C.B.L., S.M., G.H. and R.P.B. conceived the study and designed the experiments. S.M. and G.H. supervised the project. R.P.B. performed the experiments and wrote the first draft of the manuscript. All these authors reviewed the manuscript. G.B.A. and R.K provided technical support and performed bioinformatics analysis.

## Acknowledgments

We thank Judith Huirne for providing endometrial biopsies.

---

**Supplementary Fig. 1.**
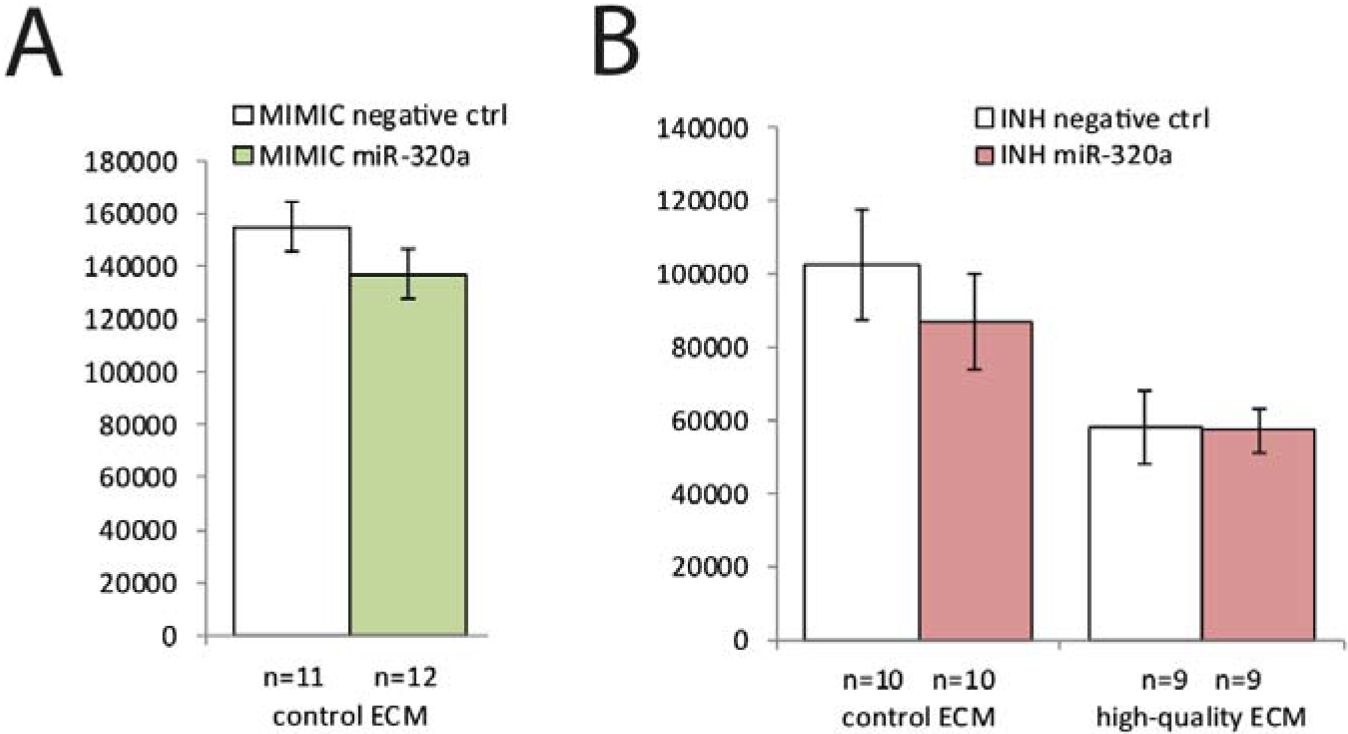
Mimics (a) or inhibitors (b) for hsa-miR-320a did not affect migration of non-decidualized hESCs when co-cultured with control ECM or high-quality ECM.

